# Chaotic dynamics in blooms of *Ostreopsis* cf. *ovata* modelled from environmental time series using ordinary differential equations systems

**DOI:** 10.64898/2026.09.25.754431

**Authors:** Andy De Almeida, Samuel Hocquart, Martin Rosalie

**Affiliations:** UMR 5096 Laboratoire Génome et Développement des Plantes, Université de Perpignan Via Domitia, CNRS

## Abstract

Blooms of the toxic dinoflagellate *Ostreopsis* cf. *ovata* can cause respiratory irritation in beach users, yet their environmental drivers and temporal dynamics remain poorly understood. Here, we use a dynamical-systems approach to investigate how environmental conditions shape bloom dynamics from long-term time series of algal abundance and abiotic variables. Using the global modelling framework GPoM, we reconstruct nonlinear ordinary differential equation systems and identify candidate models capable of reproducing key features of the observed dynamics, including chaotic behaviour. Topological and statistical analyses reduce the candidate models to two systems that best capture the observed distributions and dynamical patterns. These models reveal distinct combinations of environmental conditions associated with bloom occurrence and amplitude, providing a framework for interpreting bloom trajectories and a potential basis for short-term prediction.

## 1 Introduction

Although planktonic and aerosol measurements provide valuable information on immediate human exposure to harmful microalgae, they may not reflect the benthic cellular reservoir that sustains bloom development, where the highest cell abundances are often found (Cohu et al., 2013; Ciminiello et al., 2014). Blooms of the microalgae *Ostreopsis* cf. *ovata* have been appearing on French Atlantic coast since 2021 (Sadorge et al., 2026). They have been common around the Mediterranean for the past twenty years where it socio-economic impacts have been documented (Berdalet et al., 2022). It is necessary to understand the links between environmental variables and the occurrence of these blooms by providing models.

Recent reviews highlight the growing use of modelling approaches to describe and forecast algal bloom dynamics, with empirical and data-driven methods, including machine learning, increasingly used to capture complex and non-linear relationships between bloom dynamics and environmental drivers (Caballero et al., 2025). More broadly, current modelling approaches range from conceptual and empirical-statistical models to process-based models, with applications spanning bloom dynamics, short-term forecasting, and longer-term predictions; however, the integration and exploitation of long-term time series can improve our understanding and prediction of bloom dynamics (Wang et al., 2024).

In Italy, Accoroni *et al*. (Accoroni et al., 2015) used a dataset collected between 2007 and 2012 to investigate the environmental factors associated with the onset, maintenance, and decline of *O.* cf. *ovata* blooms. More recently, Fabri-Ruiz et al., 2024a used long-term environmental and biological observations combined with high-resolution climate simulations to model the present and future distribution and abundance of *O.* cf. *ovata* in the Western Mediterranean Sea, showing that temperature and chlorophyll dynamics were key predictors of abundance and that climate change could extend the seasonal bloom period. The environmental and biological dataset compiled and made publicly available by Fabri-Ruiz et al., 2024b provides an opportunity to further investigate the temporal dynamics of *O. cf. ovata* blooms; the analyses presented in this study are based on these recently released data in Monaco.

In this study, we aim to use their dataset (Fabri-Ruiz et al., 2024b) to investigate whether the observed dynamics of *O. cf. ovata* can be modelled by an underlying deterministic dynamical system. Rather than focusing bloom solely on statistical associations between variables, we seek to identify a system of nonlinear ordinary differential equations (ODEs) able to reproduce the main features of the observed dynamics. This approach is motivated by the possibility that apparently complex or irregular time-series patterns may emerge from relatively low-dimensional deterministic dynamics, including chaotic dynamics, which can arise from systems described by as few as three coupled differential equations (Letellier, Aguirre, et al., 2009). Identifying such dynamics would provide a mechanistic representation of the relationships among the variables and could help reveal the structure underlying the observed temporal variability.

To achieve this, we use the Generalized Polynomial Modeling (GPoM) framework, implemented in the GPoM R package, which is specifically designed to reconstruct deterministic Ordinary Differential Equations (ODE) models from observational time series and to identify low-dimensional deterministic and chaotic dynamics (Mangiarotti, Le Jean, et al., 2023). GPoM is particularly suited to our objective because it performs a systematic search for polynomial model structures and evaluates the resulting dynamical regimes, rather than requiring a predefined library of candidate functions. GPoM therefore provides a framework for our objective of exploring whether the observed algal bloom dynamics can be captured by a low-dimensional deterministic model and whether signatures of chaotic dynamics can be identified in the reconstructed system. Since its introduction in the 1990s, global modelling has proven successful in reconstructing complex dynamics across a variety of environmental systems, including cereal crop dynamics from remote sensing data (Mangiarotti, Drapeau, et al., 2014), hydrological processes (Mangiarotti, Fu, et al., 2021), and sandy shoreline–sandbar systems (Aparicio et al., 2026).

In this study, we use GPoM to generate three-dimensional nonlinear ODE systems from combinations of environmental and biological variables, focusing on models capable of exhibiting chaotic dynamics. We increased progressively the number of polynomials terms used for modelling. We then use a topological characterization of the We identify models displaying chaotic dynamics and characterised the topology of their attractor. We thus reduced the number of candidate model by retaining systems displaying distinct dynamical properties. Statistical comparisons of the model outputs and observed data are subsequently used to identify, among the remaining candidates, the models that best reproduce the observed distributions of values. This approach allows us to classify the environmental variables according to their contribution to bloom dynamics and to identify the dynamical structures associated with bloom occurrence and amplitude. Finally, we relate the template of the selected model to bloom trajectories to investigate how the structure of the underlying dynamical system may explain forthcoming bloom amplitude and occurrence, providing a basis for short-term prediction.

## 2 Material and methods

### 2.1 Material: time series

Time series used in this study have been made available as supplementary material associated to the publication (Fabri-Ruiz et al., 2024a). The dataset (Fabri-Ruiz et al., 2024b) is directly available to download at this address: https://doi.org/10.5061/dryad.44j0zpcn5. In this dataset, abundance of the microalgae in time series is part of long-term monitoring programs. Authors also provides a statistical correction of physical and biogeochemical reanalysis of the Mediterranean Sea to match in situ environmental conditions.Time series are available from 2007 to 2018 only in the summer. We used the Copernicus closest accessible data point to retrieve time series of O_2_, NO_3_, PO_4_, salinity and surface temperature seawater (Copernicus Marine Service, 2026a; Copernicus Marine Service, 2026b). The latter time series whose start from 2013 but contains seawater temperature at various depth. We fit the data of abundance including an exponential growth to reproduce observations (Pezzolesi et al., 2012) (see Fig. 8 for fit over observed points in blooms). The smoothing of the raw data provided is computed using a Savitzky–Golay filter and splines (Fig. 1). The availability of the data (2007 to 2017 for all except for temperature only available from 2013 to 2017) leads us to run the method (Sec. 2.2) on three samples of the time series: range A (2013 – 2017) to include temperature as a variable, range B (2007 – 2017) the entire data set except temperature and range C (2008 – 2016) without first and last year. In the following, we mostly discuss A (with temperature) and C (without temperature) because model selection based on sample B was highly driven by the initial flowering of 2007 and with less confidence in the fitting of the original data with our curve for the last year (Fig. 8).

**Figure 1:**
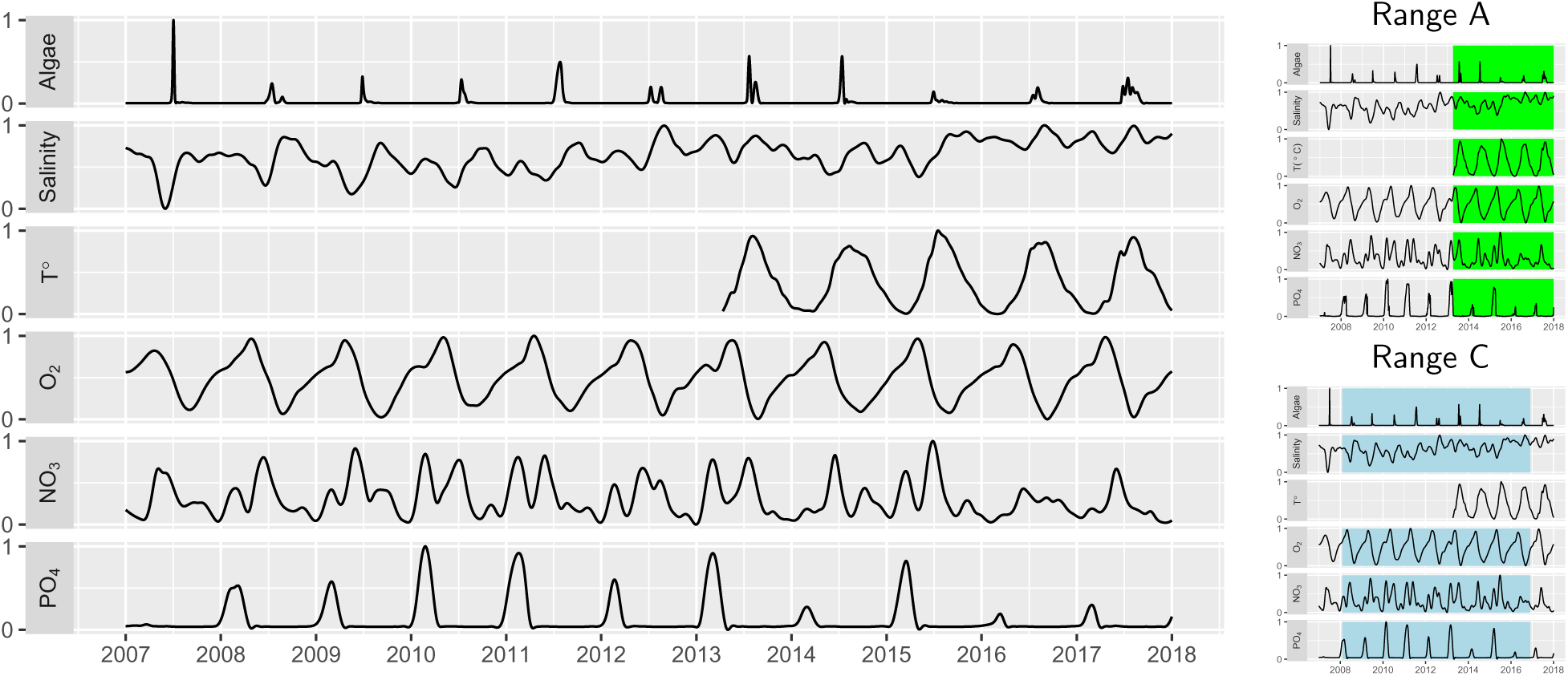
Time series of five variables that are considered of interest to impact blooms. Variables have been extracted from the dataset provided in the original publication (Fabri-Ruiz et al., 2024a; Fabri-Ruiz et al., 2024b) from Monaco. The original variables are normalized so that their values lie between 0 and 1. Sample used for modelling are indicated on the side plot by underlying the range of values considered for modelling.

### 2.2 Methods

#### Global polynomial modelling

The GPoM framework (Mangiarotti, 2015) automatically generates candidate polynomial dynamical models (Ordinary Differential Equations systems), estimates their parameters from the time series, and selects the model that provides the best fit according to predefined selection criteria. In a typical use of GPoM, one can specify a range for the number of polynomial terms in the models, as well as the maximum degree of the polynomials (terms or combinations of terms, fixed here to 3). In this study, we propose to gradually add new polynomial terms to introduce greater complexity into the generated models (from 5 to 30 polynomial terms in). Our GPoM modelling process is designed to look for deterministic relationships between tested variables by generating three dimensional ODE systems from three time series: algae and two other variables. The GPoM package is an open-source package developed in R language and made available on the Comprehensive R Archive Network (CRAN) (Mangiarotti, Le Jean, et al., 2023).

#### Topological characterisation

Topological characterisation method provides the structure of chaotic attractors to underline the spatial transformation of the trajectories by establishing a template. The procedure is described with some steps Fig. 2 for the standard Rössler attractor (Rössler, 1976) as an example (results are originally produced in (Letellier, Dutertre, et al., 1995)). This attractor (Fig. 2A) is bounded by a genus–1 torus, thus a Poincaré section with only one component is required. For attractors bounded by higher genus torus, the number of components varies (Tsankov and Gilmore, 2004) inducing changes in the template structure (Rosalie and Letellier, 2014). The Poincaré section is the crossing of the trajectory to a plane. and, from it, a variable *ρ_n_* is defined to represent the distance from the inside of the attractor to the outside. This variable is used to build the *first return map* that is the signature of the chaotic dynamics (Fig. 2B). This topological characterisation method relies on the linking number between these UPOs (Fig. 2A), this topological invariant quantifies the relative rotation rates of these UPOs considering them as mathematical links. The template (Fig. 2C) syntheses the three main properties of chaotic attractors: (i) the *global time invariant flow* from a deterministic model are represented with a bounded structure, (ii) the *sensitivity to initial conditions* is visualised with stretching, tearing, folding and squeezing mechanisms and, finally, (iii) the *aperiodicity of the solution* can be underlined by the unstable periodic orbits contained in the template that are approached but unreached during the solving of the system. The validation of a template is obtained by having the theoretical linking number from the template equals to those obtained numerically. We developed a tool to perform the first steps of this methodology automatically to accelerate detection of similar dynamics (Hocquart and Rosalie, 2026).

**Figure 2:**
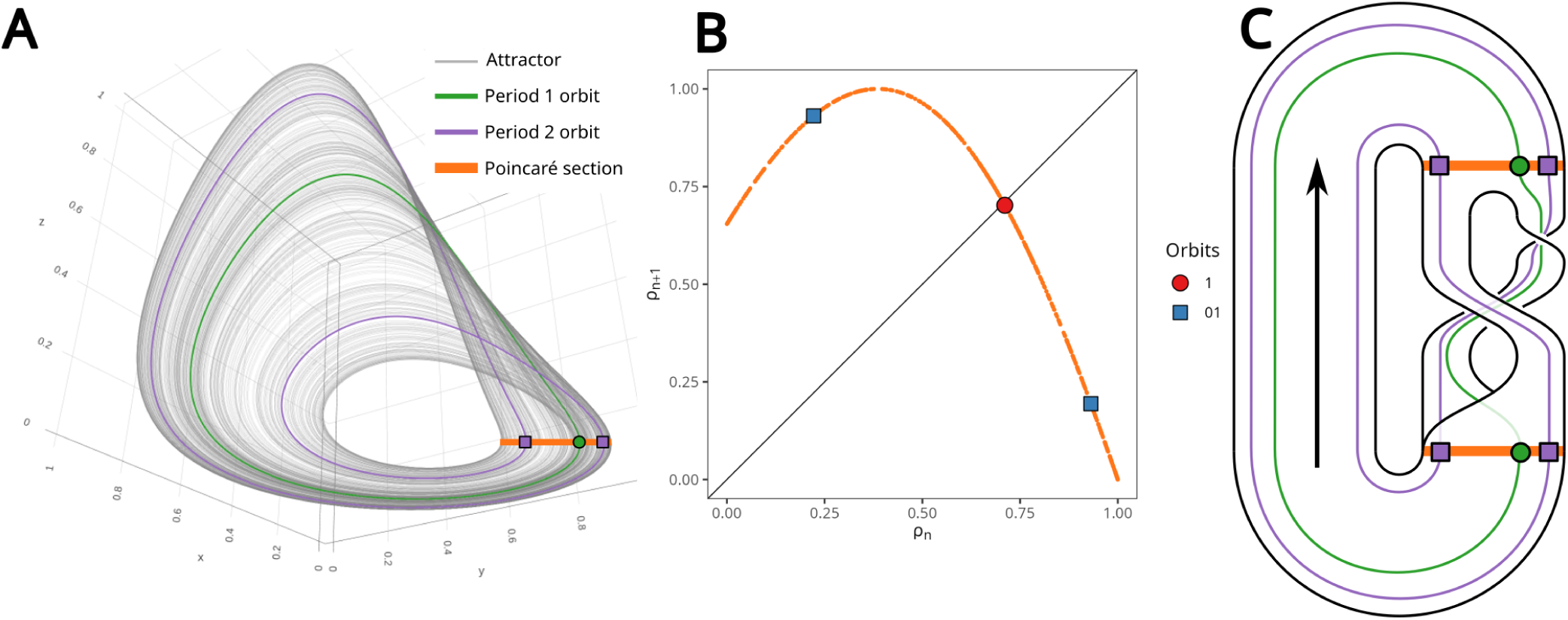
Steps of the topological characterisation of a chaotic attractor. A. Chaotic attractor solution of the Rössler system (Rössler, 1976) in a three-dimensional space. The orange line is the Poincaré section and with this projection, the flow evolves clockwise. From the Poincaré section, the Unstable Periodic Orbits (UPOs) are extracted numerically (for example, two orbits are ploted). B. First return map to the Poincaré section with periodic points (UPOs crossing the Poincaré section). The structure with an increasing branch followed by a decreasing branch directly indicates the number of branches in the template. C. Template describing the topological properties of the attractor. A template is validated when theoretical linking numbers between orbits (computed from the template) correspond to thoose obtained numerically (A).

For attractors bounded by a genus–1 torus, the foliated structure with additional branches in the first return map is a delay before squeezing (Meneceur et al., 2025): there is no need to distinguish them as extra branches. Oppositely, for genus–3 bounded torus, the foliated structure must be explicitly detailed in the template (Rosalie, 2016). Obtaining a topological template becomes difficult for attractors bounded by a high genus torus and a finely foliated structure (Rosalie, 2016; Tabekoueng Njitacke et al., 2025; Letellier, 2026).

#### Model selection

From the set of models generated by our GPoM modelling process, numerous three-dimensional ODE systems are generated to propose models linking algae dynamics to two other variables. GPoM tool already include a classification procedure but some particular dynamics (such as transient chaos) are not easily distinguishable from stable chaotic dynamics. The first step is to determine the dynamics of the GPoM “unclassified models”: converge, diverge, stable oscillations or chaotic dynamics to reduce the number of candidate models.

First-return maps — and templates, when available — serve to prune the pool of candidate models. When multiple candidates yield identical templates, the principle of parsimony is applied to select the model with the fewest terms. In the absence of a discernible template, the geometric structure of the first-return map is combined with the linking numbers of unstable periodic orbits to evaluate the dynamical equivalence between systems.

The comparison between the distribution of the empirical data used for modelling and that of the data generated from the systems’ numerical simulations ensures that the model accurately reproduces the observed statistical distribution. This comparison is conducted using the Kolmogorov-Smirnov test (Massey, 1951), a non-parametric statistical method that quantifies the distance between the cumulative distribution functions of two samples to assess whether they originate from the same underlying distribution.

The identification of these models implies an underlying determinism in the summer blooms. This determinism is essential, as it constitutes one of the fundamental properties that must be verified before establishing the presence of chaotic dynamics. Regarding the structure of the proliferation peaks, the geometry of the attractor — specifically whether it is dissipative or weakly dissipative — directly impacts the structure of the modelled summer blooms.

## 3 Results

### 3.1 Variable selection

Fig. 3 displays the number of unclassified models generated by GPoM from the time series. To illustrate the implications of these results, we examine two specific cases Fig. 3C. First, fewer than twenty ODE systems coupling microalgae with O_2_ and PO_4_ are identified, regardless of the maximum number of terms allowed in the model. In contrast, it highlights that 98 ODE systems link microalgae dynamics with salinity and NO_3_ when the model is restricted to 25 polynomial terms (*x* = 25). For this latter pair of variables, a linear trend emerges, indicating that increasing the number of permissible polynomial terms leads to a proportional increase in candidate unclassified dynamical systems.

**Figure 3:**
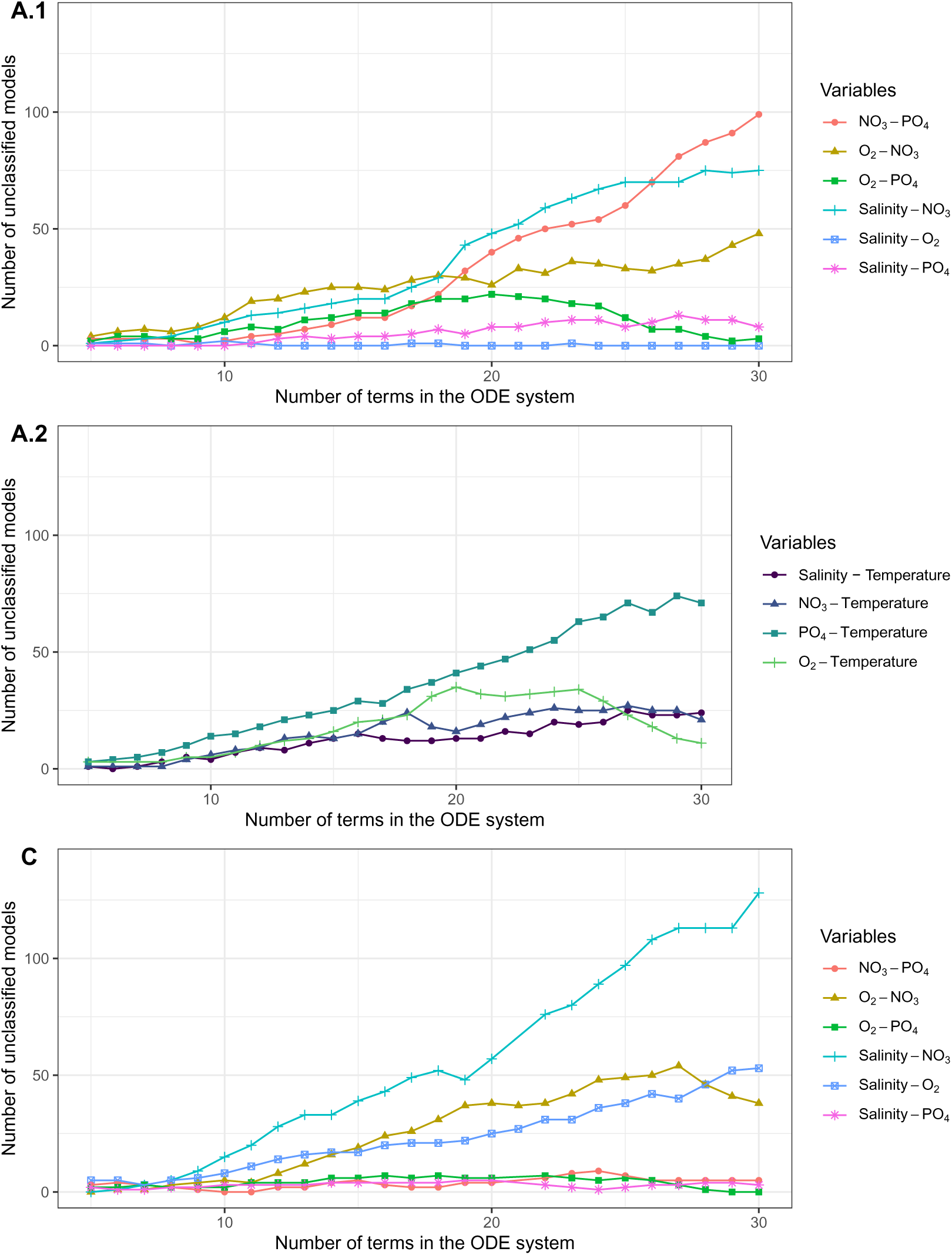
Number of unclassified systems generated by GPoM for couple of variables. The *x*-axis value is one of the parameter used as an input of GPoM::gpomo method in R (Mangiarotti, Le Jean, et al., 2023). Produced systems are filtered to avoid proposing systems that diverge or converge to a point; others reported here are considered as unclassified. Experiences A and C (associated to respective plots labels A.1, A.2 and C) differ from the portion of time series used to generate models (Fig. 1).

From a broader perspective, across all panels in Fig. 3, it is noteworthy that regardless of the selected time-series segment, the graphs consistently indicate a vast number of models coupling salinity and nitrate. Given that identifying a valid model from purely random data is virtually impossible (Letellier, Aguirre, et al., 2009), the systematic extraction of these equations underscores a strong dynamical link between these variables. This coupling is further emphasised by the contrast with other variable pairs, such as phosphate and oxygen, for which far fewer unclassified models are retrieved. Fig. 3A.1 underline the coupling with temperature, algae and another variable and less than 40 models are retrieved by most of them, where a plateau is reached when more than 20 terms are allowed in ODE systems. Only the couple temperature with phosphate seems to follow the linear trend indicating that this association could be an option for modelling the blooms.

### 3.2 Dynamics and topological characterisation

Fig. 9 is the numerical integration of the 1600 unclassified ODE systems generated for three couples of variables (salinity - NO_3_, salinity - O_2_ and NO_3_-O_2_) in Fig. 3C. Long numerical integration of these systems is required to avoid having systems with transient chaos which ultimately converge or diverge. An example with times series and attractor is provided, Fig. 10, to illustrate an example of classification measures performed. It indicates that half of the unclassified systems diverge, while 12% converge to a point and 36% converge to a limit cycle. Only 2% of these systems converge to a chaotic attractor and they are all linking microalgae blooms with salinity and nitrate. This type of model with chaotic dynamic seems to successfully capture the dynamics by providing us ODE models able to have the variety of blooms observed in the time series that the others (divergence, convergence or limits cycle) cannot. The dynamics of these chaotic solutions are studied in the following subsection to compare the systems and characterise their structure.

The Fig. 4 displays the two key components used to categorize and compare the dynamics for each of the three preceding experiments (Fig. 3). The genus of the bounding torus provides insight into the flow dynamics. A genus-1 torus indicates that the surface undergoes only stretching and folding before returning to the Poincaré section; this restricts the range of distinct dynamics and reveals numerous equivalences among certain systems. Conversely, a genus-3 torus implies that the flow is torn apart, thereby increasing the complexity of the dynamics.

**Figure 4:**
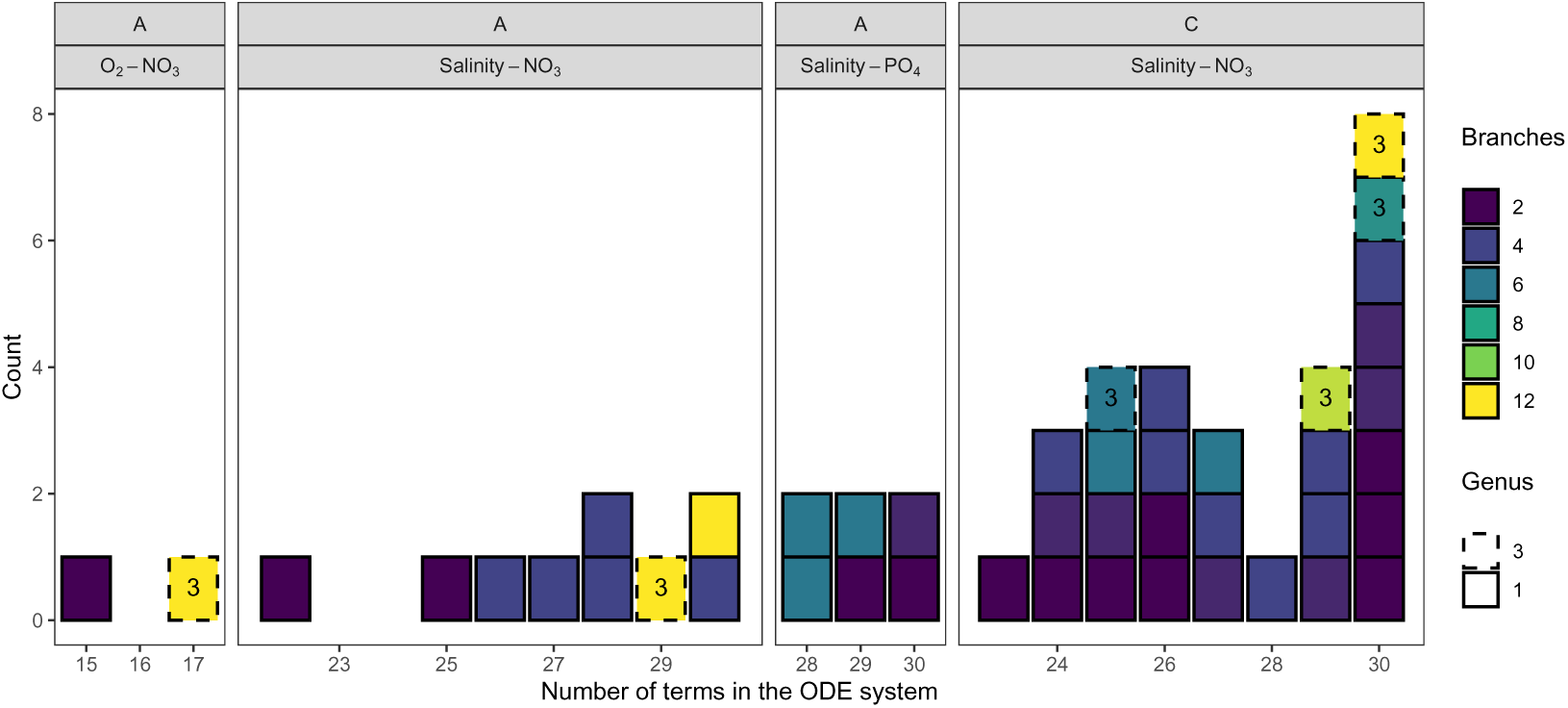
Structure of chaotic attractors solution of ODE systems by bounding torus and number of branches. Distribution of structural values for chaotic dynamics solution to the systems of Fig. 3. The genus of the bounding torus gives the global architecture of the templates allowing higher complexity. The number of branches indicates the main pathway the trajectory can follow in scenarios until the next topological period.

The second parameter corresponds to the number of branches identified in the first-return maps. As demonstrated in the appendix (Figs. 11, 13 and 14), this configuration implies numerous possibilities regarding the system’s scenarios, given that the first-return map and its corresponding topological template share the same number of branches to characterize the dynamics. Consequently, the template serves as a global representation of all admissible evolutionary pathways. The branch line of the template, preceded by stretching and squeezing mechanisms, indicates the transition between distinct dynamical regimes, which accounts for the high diversity of potential trajectories and, thus, scenarios for amplitude of algae blooms.

However, templates could not be reconstructed for every attractor compiled in Tab 3. The stationarity status of the time series (Tab. 2), especially salinity, induces foliated structure (attractor with multiple similar layers in the phase space) in most of the models generated as it has been reported theoretically (Aguirre and Letellier, 2012). Nonetheless, the first-return map provides adequate insights to categorise the systems and select those exhibiting the highest dynamical complexity. Foliated structure for attractors bounded by a genus–1 torus induces structural repetitions (models 11 and 27 of Tab. 3) that do not occur for attractors bounded by genus–3 torus due to their structure. For example, the template of attractor 40 of Tab. 3 that can be visualised Fig. 14 while advances tools are required to obtain the template of attracor solution of model 9 of Tab. 3 (Fig. 12). Appendix Tab. 5 presents the first four statistical moments of two models with distinct dynamics that cannot be inferred from the statistics alone.

**Table 1:** The similarity between the distributions was assessed using the two-sample Kolmogorov– Smirnov test, with the Kolmogorov–Smirnov D statistic (*D*) used as a measure of distributional distance. Entire time series is used to made the comparison although, depending on the assumptions (range A, B or C), only a portion was used to generate the models.

| | Variable | $D$ | $p$-value |
| --- | --- | --- | --- |
| Model 2 | Algae | 0.8541219 | 0 |
|  | O<sub>2</sub> | 0.1296102 | 4.976814e-96 |
|  | NO<sub>3</sub> | 0.2144508 | 3.722993e-262 |
| Model 9 | Algae | 0.55731 | 0 |
|  | Salinity | 0.36242 | 0 |
|  | NO<sub>3</sub> | 0.36242 | 0 |
| Model 11 | Algae | 0.70641 |  |
|  | Salinity | 0.25377 | 0 |
|  | NO<sub>3</sub> | 0.33033 | 0 |
| Model 12 | Algae | 0.89501 | 0 |
|  | Salinity | 0.84012 | 0 |
|  | PO<sub>4</sub> | 0.48333 | 0 |
| Model 27 | Algae | 0.41796 | 0 |
|  | Salinity | 0.10972 | 6.196963e-69 |
|  | NO<sub>3</sub> | 0.47484 | 0 |
| Model 29 | Algae | 0.540449 | 0 |
|  | Salinity | 0.15796 | 2.01877e-142 |
|  | NO<sub>3</sub> | 0.378696 | 0 |

Finally, as described in section 2, Tab. 3 details the pruning of the 48 models of (Fig. 4) by listing their genus and number of branches in templates. At least, one selected model per set of three columns: sample dataset, variable and genus, except when there is only a two branches template bounded by a genus–1 torus. This pruning leads to a set of eight models (Tab. 3): one for A - O_2_ - NO_3_ (genus–3 only), two for A - salinity - NO_3_ (genus–1 and genus–3), one for A - salinity - PO_4_, one for B - O_2_ - NO_3_, one for B - PO_4_ - NO_3_ and two for C - salinity - NO_3_ (genus–1 and genus–3). However, moment statistics (Tab. 4) indicate that models B - O_2_ - NO_3_ and B - PO_4_ - NO_3_ are out of range in term of values, we do not select them, six models remain with plausible moment statistics.

### 3.3 Model selection and validation

An examination of the phase space and variable distributions inform on the structural characteristics of the models and evaluates their alignment with the empirical data (Fig. 5). It should be noted that in this representation, the entire time series used is shown across the whole visualisation, even though this series is truncated for hypotheses A and C. For instance, in the third line (model 11), the highest peak of algae is not part of the data used for modelling as it is the first bloom recorded in 2007 and in hypothesis A, values selected from 2013 to 2017. A similar rationale applies to the statistical comparisons using the Kolmogorov–Smirnov test (Table 1); the objective is to perform a pairwise model comparison regardless of the specific hypotheses used for their generation.

**Figure 5:**
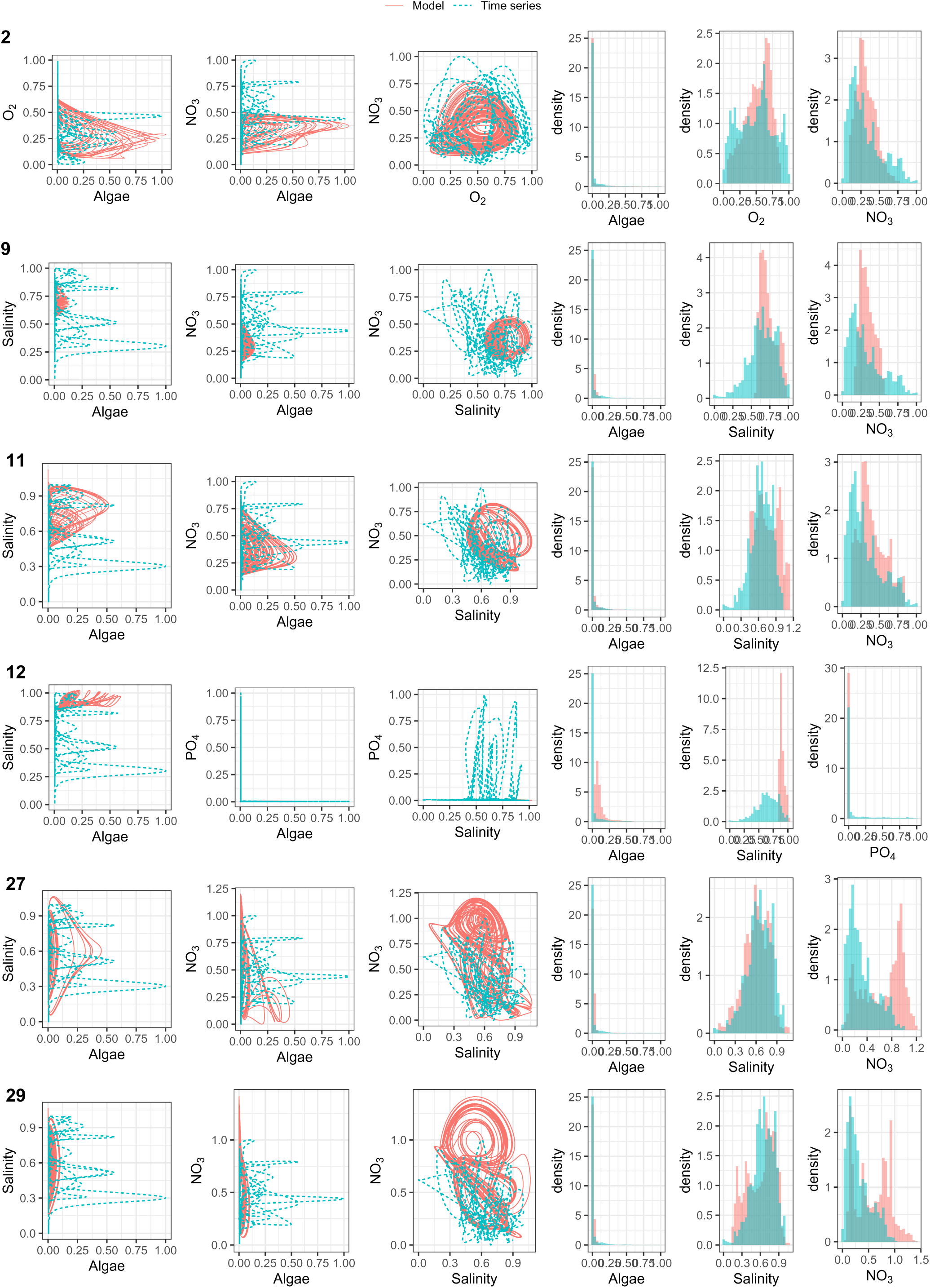
Phase space and distribution of values of the models. One row per model, correspondence of the number of the model is in Tab. 3.

The model with the lowest *D* on algae and salinity (Tab. 1) is the model 27 with initial conditions *A*(*t* = 0) = 0.000445281, *S*(*t* = 0) = 0.804196 and NO_3_(*t* = 0) = 0.553187:

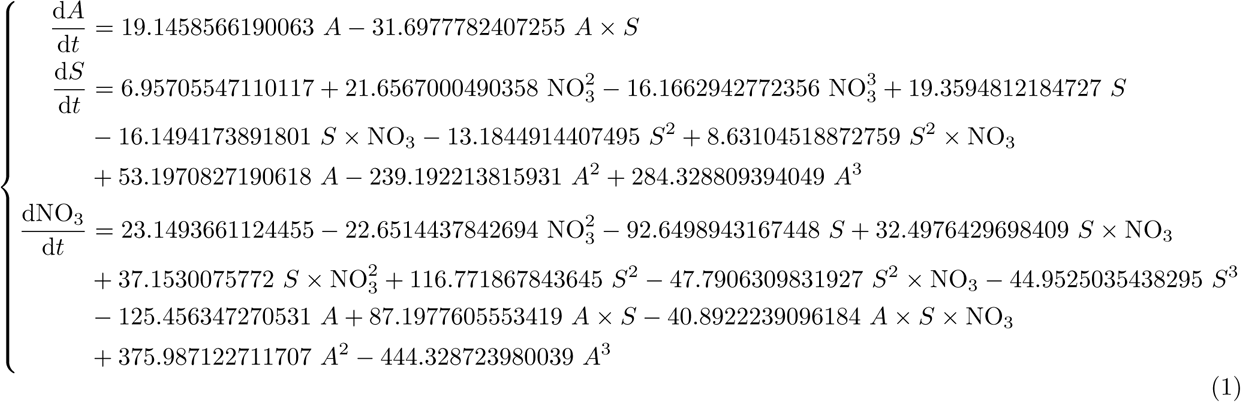

The distributions (Fig. 5) illustrates that first model choice and for the third variable (NO_3_), the value *D* is one of the highest but, the distribution captures the initial peak of the times series and emphasises the peak appearing in the distribution of the time series. This model is best suited to representing the joint variation of these variables (Algae – Salinity – NO_3_).

Model 2, for its part, is of interest because, although it was trained on only the final half of the data, the distances for salinity and oxygen are relatively low compared with the other values and therefore deserve our attention. Model 2 is defined as follows with initial conditions: *A*(*t* = 0) = 2.2789.10*^−^*_12_, O_2_(*t* = 0) = 0.565424 and NO_3_(*t* = 0) = 0.253898:

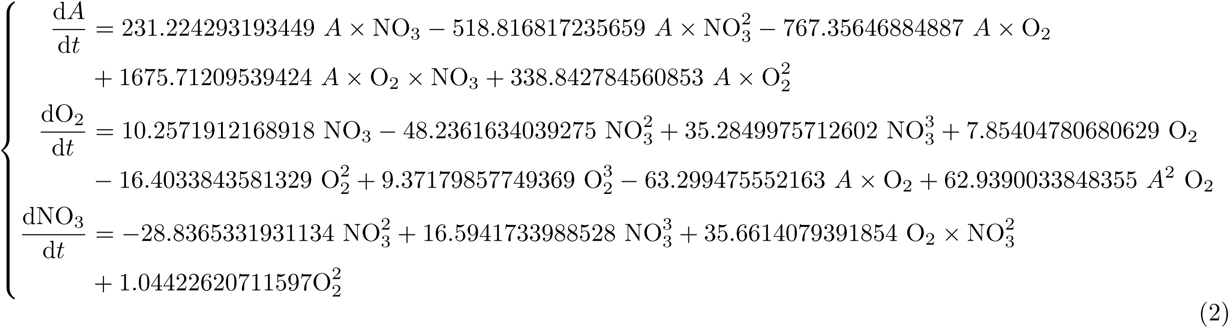

Although microalgae blooms are of significant interest, their onset and magnitude cannot be fully explained by current models; hence, we seek to characterise and model these phenomena. In the models, these blooms are represented Fig. 6 while original blooms are in Fig. 8. Both models have the capacity to reproduce this structure with values close to 0 for a long period before the bloom. Their structures differ because model 27 can have oscillation before peak with a lower amplitude. This is not observed in model 2 but it produces blooms with a wider range between each. From a statistical standpoint, these models were retained due to their consistent value distributions; furthermore, they successfully replicate the dynamic behaviour of the blooms.

**Figure 6:**
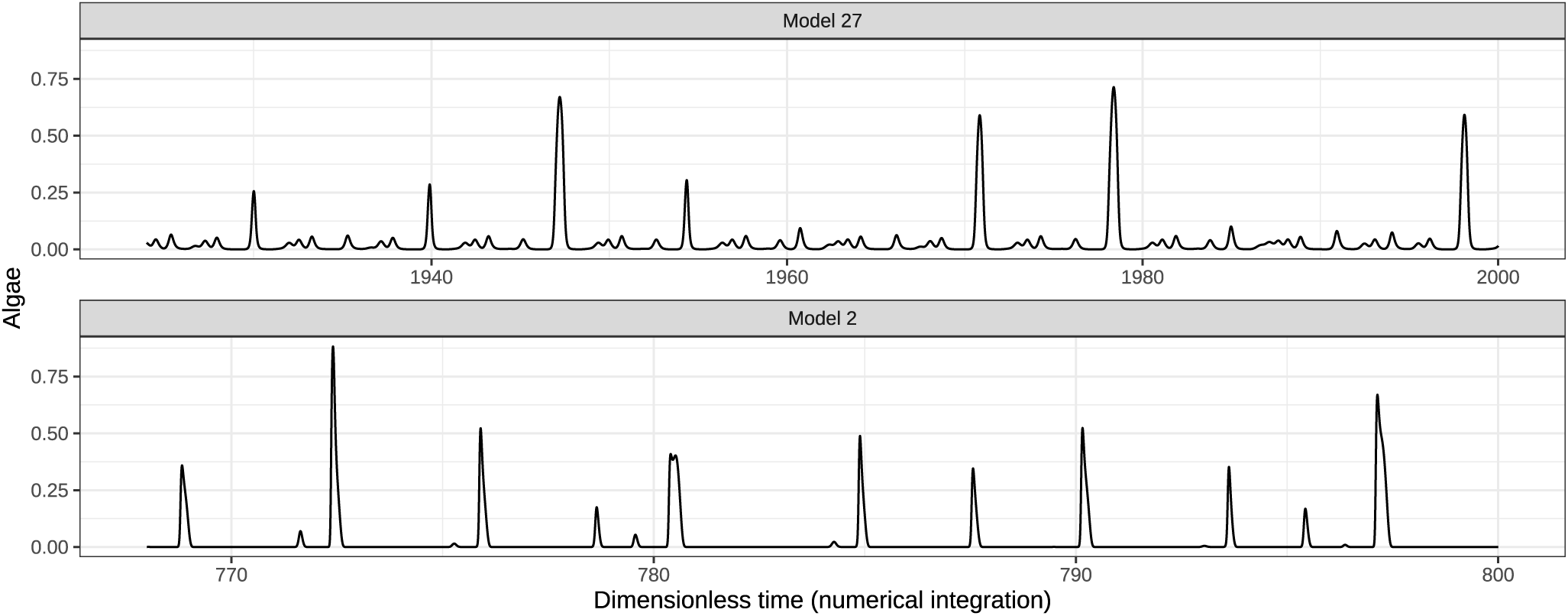
Time series and blooms produced by models 27 (salinity and NO_3_) and 2 (O_2_ and NO_3_). The values are the result of a numerical integration of the two models (1) and (2).

## 4 Discussion

### 4.1 Selection of variables in models

Regardless of sample dataset use to select the models, the models incorporating salinity and NO_3_ are more prone to display a chaotic dynamics able to reproduce algal variations in abundance. Increasing the number of polynomials permitted in the ordinary differential equations models leads to more complex dynamics, which better reproduce the various blooms (Fig. 4). The fact that deterministic models have been obtained is a strong indicator of the relationships between these variables. This is even more so given that it does not appear to be such a direct link between algal blooms and phosphate or temperature according to our numerical experiments.

By definition in ordinary differential equations, derivatives are intrinsically embedded in the model structure; hence, they are not treated as separate categories in our classification contrary to the original publication (Fabri-Ruiz et al., 2024a). Consequently, the number of unclassified systems generated from GPoM in Fig. 3 indicates that couple NO_3_ and salinity comes first in proportion, followed by the pair temperature with PO_4_. From a dynamical perspective, the generated models including temperature only yield limit cycles, or solutions that either converge or diverge. This does not mean that temperature has no influence, but there are no triplet of coupled variables including temperature and algae abundance that can generate chaotic dynamics. The surface temperature has been selected for these experiments but the temperature of other layers of seawater should have been more appropriate, moreover, these data are also available on Copernicus (Escudier et al., 2021). Previous studies have identified water temperature as the primary driver of *O.* cf. *ovata* dynamics, with temperature largely defining its ecological niche (Cohu et al., 2013; Carnicer et al., 2015). In contrast, our results suggest that temperature is not the main environmental factor explaining the observed patterns in our study system, While temperature is certainly a major driver of *O. ovata* dynamic, it is probably not selected because temperature is not influenced by the other variables. In contrast, we can find feedbacks between the concentration of algae and the concentration of phosphate, nitrate and O_2_, that are consumed and/or released during *O. ovata* blooms. Our recurrent finding that salinity is onemain variable coupled with algal concentration is even more surprising: one does not except salinity to be influenced by any of the presented variables, except temperature. A strong positive correlation exists between the accumulation of positive spring sea surface temperature anomalies and the onset date of the blooms, suggesting that higher spring temperatures promote earlier *O.* cf. *ovata* blooms in Monaco (Drouet, 2020). This implies that temperature primarily regulates the temporal onset of the phenomenon rather than its magnitude or structure. Overall, the impact of temperature is more complex, given the varying observations of blooms at different times of the year (Drouet et al., 2022).

From the Fig. 4 combined with selected model (Fig. 5 illustrating all chaotic dynamics structure, it can be inferred that salinity and NO_3_ should initially hold equal prominence within the model structure. Concurrently, decreased salinity has been shown to inhibit *O.* cf. *ovata* growth, with the highest growth rates reported under relatively high salinity conditions (Pezzolesi et al., 2012). This is consistent with our results, which highlight salinity as an important driver of *O.* cf. *ovata* abundance. As for nutrients, they obviously have an impact on the growth of this microalgae (see Vanucci et al., 2012 for details). Thereafter, O_2_ followed by PO_4_ may be integrated as supplementary factors to effectively forecast bloom events.

Regarding PO_4_, a temporal lag between the two variables may also need to be considered, as elevated PO_4_ concentrations tend to coincide with lower values of the other variable. Delay differential equations (DDEs) could therefore provide an alternative modelling framework for explicitly accounting for such temporal dependencies and delayed effects. Some algorithms have been developed to generate this type of equations from time series (Breda et al., 2026) and it can be a complement to this work with GPoM.

### 4.2 Dynamical properties of models

For a third of the system reported in Fig. 4, the dynamical properties exhibit the simplest chaotic mechanism with only two branches. Only 12 systems (25%) has dynamics with more than four branches. Within the same couple of variables for a model of Fig. 4, the dynamics remain close to one another; adding polynomial terms therefore increases the dynamical complexity without changing its effective coverage on the data. However, the genus of the bounding torus of an attractor has an impact of the dynamical structure of attractors and consequences the predicted dynamics. The complex dynamics of attractors bounded by genus-3 (or higher) tori indicate that, for close initial conditions, the stretching mechanism causes the trajectories to split into two separate directions: the flow divides. This contrasts with mechanisms solely based on stretching and folding, which exhibit a relative continuity although the trajectories continue to diverge exponentially. This means that some of the identified models exhibit this property (genus–3 bounded torus), which could imply that, despite starting from very similar initial conditions, they may nonetheless produce substantially different short-term dynamics. Under such dependence to the initial condition, one expect that correlative studies using static data or short term time series would not find the mechanistic link between environmental parameters and algal growth, which alter their predictive value.

Analysis of the models’ dynamics revealed that the data induce a foliated structure. This foliation leads to repeated structural patterns that may resemble regular, damped or amplified oscillatory motifs observable in the time series (Fig. 8 and Fig. 6). This reflects a chaotic dynamic characterised by successive stretching and folding, for which collapse occurs only after traversing multiple foliations. This occurs because the trajectory is deterministic and requires multiple cycles to reach the squeezing mechanism, which is synonymous with the mixing of trajectories. All these distinctive dynamics were captured by the different models, and the subsequent selection process identified the model that best reproduces dynamics of the data.

The model we propose cannot be directly interpreted in biological terms, given the nature of the polynomial terms composing its equations. Nevertheless, it provides insights into the relationships linking these variables. Following the approach adopted by Aparicio et al., 2026, one could investigate whether only a subset of these terms predominates during the numerical solution and thereby reduce the complexity of the model. Alternatively, complexity could be reduced by identifying a simpler system of differential equations that is algebraically more tractable while remaining topologically equivalent and reproducing the dynamics of Model 27.

### 4.3 Chaos and template: towards scenarios for blooms

Combining parcimony principle with topological characterisation of chaotic attractors provides information regarding dynamics. Among the most numerous models and the one ultimately selected, couple salinity and NO_3_, there are strong positive correlations between the number of models generated and the fixed number of polynomials (Fig. 3). This trend is also reflected in the dynamic complexity of the generated systems (Fig. 4), which shows that the tool can produce models capable of better capturing the dynamics of the data when used in this way. The drawback is that a significant number of dynamic systems must then be analysed (13,864 for this study, based on three hypotheses) where 48 chaotic attractors has been identified.

By intrinsic definition chaotic dynamics are not predictable (sensitivity to initial conditions), however, other properties allow to define scenario via templates and give range of values leading to peak amplitude categories. However, by comparing the performance of models (Tab. 1) on the entire time series even if they have been obtained on various portions (hypothesis A, B and C), we ensure that our model is validated over the entire data set.

The templates obtained for some of the attractors can be interpreted as sets of possible scenarios, with the permitted transitions representing potential short-term future trajectories. For model 27 (Fig. 7), the template is made of six branches indicating that for each topological period, the trajectories will start from the Poincaré section where the trajectory is split in six directions. As the process is deterministic, the trajectory cannot escape this path until it reaches the end of the template where all branches are stretched and squeezed to the branch line. Depending on the trajectory laid here, the next branch will be followed until the next topological period. Thus, this template details the six evolution scenarios of the trajectory. It can be viewed as restriction path for a short period to predict the trajectory evolution and consequently blooms amplitude (Fig. 7).

**Figure 7:**
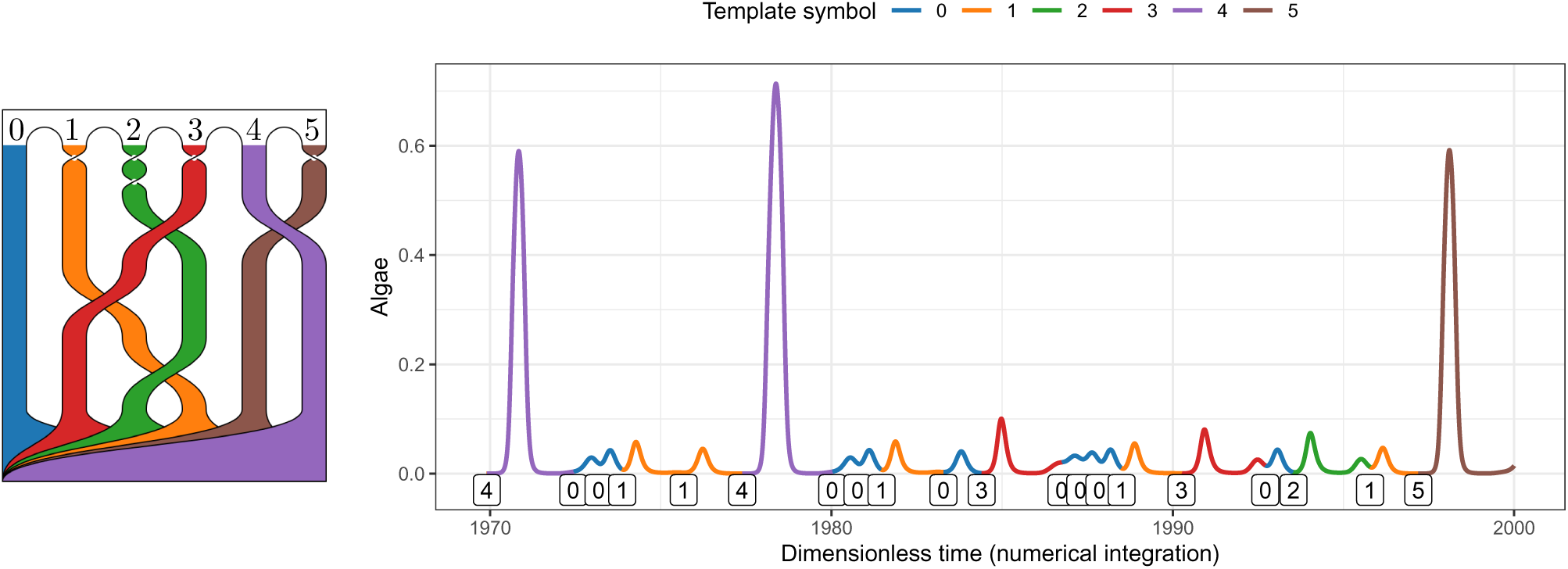
Template and time series encoding for model 27. Symbols of the template of the attractor encode the time series. Colours of the branches of the template are associated to the blooms in time series indicating the type of blooms with a distinct range of amplitude and a specific lead time before it occurs.

**Figure 8:**
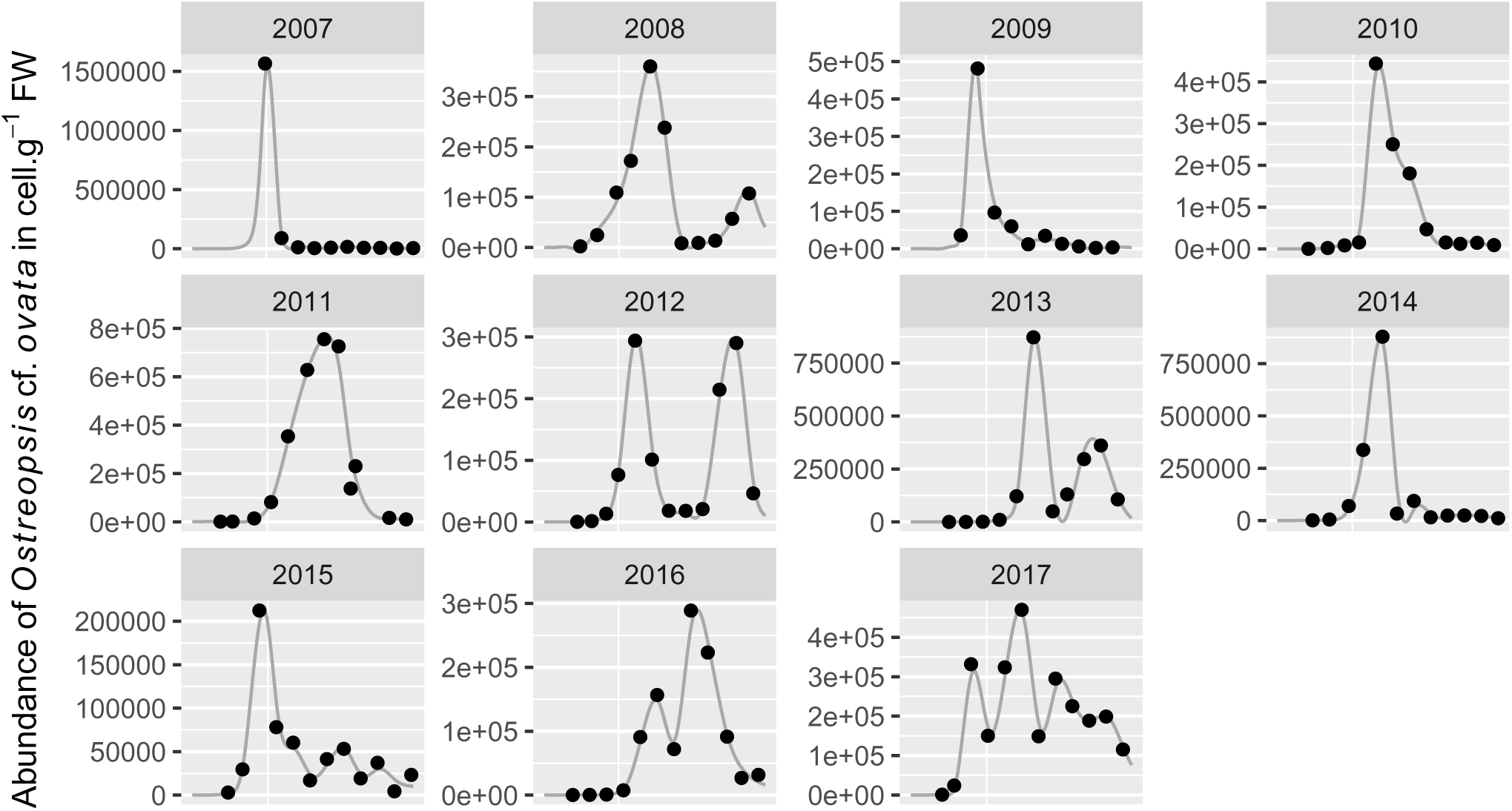
Algae abundance: original data (points) and smooth version (line). The original data come from the dataset of Fabri-Ruiz et al., 2024a (Fabri-Ruiz et al., 2024b) where the abundance unit is cells.g^-1^ FW (the number of cells per gram of fresh weight of macroalgae). Values of microalgae *O* cf. *Ovata* have been collected in Monaco. The growth rate of the microalgae is used as a biological parameter to estimate the structure of the exponential growth during the blooming period with a value of 1.55 per day. The smooth values are obtained with the method smooth.spline.

Although some of the attractors generated using GPoM could not be topologically characterised, the construction of multi-component first-return maps (Fig. 12) provides a means of more effectively tracking the evolution of trajectories (Rosalie and Letellier, 2015; Rosalie, 2016). Ideally, obtaining the template of model 2 would have enabled a direct comparison of its dynamics with those of model 27; however, this was not possible because of the complexity of its structure, which limits our ability to fully characterise and interpret its dynamics. Tools such as coloured tracer (Rosalie and Mangiarotti, 2025) or homologies with templex (Charó et al., 2022) could help to describe its structure.

## 5 Conclusion

The precise link between *Ostreopsis* cf. *ovata* outbreaks and the toxicity of sea spray is under investigation to clearly formalise the biological process. Using time series, we have generated multiple models of ordinary differential equations linking the abundance of these microalgae to environmental data. The aim of our work is to identify the factors determining bloom dynamics by investigating whether there are deterministic solutions that would explain their occurrence and magnitude. By definition, non-linear systems are good candidates for the search for pseudo-periodic event and non-regular blooms by amplitude and occurrences. With determinism and additional properties satisfied here, chaotic dynamics has be observed as solution of generated models.

We first demonstrate that the global modelling tool (GPoM) successfully generates 48 deterministic models having chaotic dynamics as solution. Most of these systems link blooms of *O.* cf. *ovata* with salinity and nitrate. We carried out a topological characterisation of most of these systems to compare the dynamic structures they exhibit. We then reduced the number of models capable of explaining the blooms by retaining only the most parsimonious models with the highest number of branches in their dynamics. The bounding torus is another factor we considered because it involves more significant structural changes in the dynamics. A statistical test comparing the data generated by the model with the time series enabled us to select one model linking salinity and nitrates to algal blooms, and another linking oxygen levels and nitrates.

By having three hypothesis (A, B and C) the tool demonstrates his robustness to our deduction by providing a large number of models linking blooms to salinity and NO_3_. With such deterministic models that predict algae population peaks, we could use the average concentrations (Salinity - NO_3_) or (O_2_ -NO_3_) prior to the peaks to make forecasts. Considering variables importance, we conclude that salinity and nitrate are the main factors, thus oxygen and phosphate could complement models. Our study highlights the fact that water temperature does not appear to be a major factor in the occurrence of algal blooms, even though it does have an impact on the other environmental variables we studied.

The models selected describe the dynamics and conditions of emerging bloom by providing a deterministic process linking the evolution of the variables. The properties of chaotic dynamics impose that prediction from collected data is not a goal due to sensitivity to initial conditions. However, the topological characterisation of the dynamics gives scenarios with short-terms pathways trajectories will follow. Finally, the bounding torus of attractors generated by five models is high and obtaining their template is a complex task requiring additional investigations to compare the dynamics. This high bounding torus implies that closed trajectories will split into two distinct directions at a point by tearing. In time series, it can be viewed as similar trajectory that cannot be dissociated before following two distinct paths with highly different type of blooms. This can be interpreted as an illustration of the intrinsic complexity of forecasting such bloom events.

These data where collected ten years ago and for future works, we plan to find recent data to first compare the dynamics with current dynamics and secondly, to first compare with the results presented here and then establish current models. Although we did not obtain directly interpretable models, adopting the technique employed by (Aparicio et al., 2026) would allow us to identify the key components within the selected model. By evaluating the contribution of each term throughout the numerical simulation process, we could determine whether these mathematical structures correspond to known biological processes. An alternative direction for future research based on this chaotic model involves imposing a forcing on the other model variables (Salinity and NO_3_) to study the time-dependent evolution of the microalgae quantity as it has been done for satellite observations and meteorological predictions to predict CO_2_ evolution in Altamira cave (Sáez et al., 2024). It may also be pertinent to investigate potential higher-dimensional models involving four or five variables, even though the three-dimensional formulations already demonstrate a capacity to reproduce both the summer blooms and complex dynamics.

## Acknowledgments

Authors thanks the University of Perpignan Via Domitia for financial support via the UPVD-BQR-2026-GP11 (*Bonus Qualité Recherche*). This study is set within the framework of the “Laboratoires d’Excellences (LABEX)” TULIP (ANR-10-LABX-41) and of the “École Universitaire de Recherche (EUR)” TULIP-GS (ANR-18-EURE-0019). The authors thank Marie Rescan for helpful remarks and comments.

## Author contributions

Conceptualisation: MR. Methodology: MR. Investigation: ADA, SH, MR. Visualisation: SH, MR. Funding acquisition: MR. Supervision: MR. Writing-original draft: MR. Writing-review & editing: All authors.

## Data and materials availability

All data needed to evaluate the conclusions in the paper are present in the main text and/or the Supplementary Materials.

## Supplementary Materials

### A Original time series

**Table 2:** Stationary tests on smoothed and normalised data (Fig. 1). Bold figures indicate that, according to this statistical test, the time series is stationary.

| | KPSS<br>$p - \text{value} > 0.05$ | PP<br>$p - \text{value} < 0.05$ | ADF<br>$p - \text{value} < 0.05$ |
| --- | --- | --- | --- |
| Algae | **0.1** | **0.01** | **0.01** |
| Salinity | 0.01 | 0.573 | **0.01** |
| Temperature | **0.0528** | 0.976 | **0.01** |
| O<sub>2</sub> | 0.0232 | 0.828 | **0.01** |
| NO<sub>3</sub> | 0.01 | 0.297 | **0.01** |
| PO<sub>4</sub> | 0.01 | **0.01** | **0.01** |

### B Dynamics of systems

A total of 13,864 ODE systems have been generated, and they are “unclassified” by GPoM (Fig. 3): 36% for hypothesis A, 43% for hypothesis B and 21% for hypothesis C. Metric tools such as Lyapunov exponent can be used automatically on the numerical solution but as the number and the variety of systems is huge, a reproducible algorithm has not been used here. We analyse a set of plots to perform the classification of the dynamics (Fig. 10) and chaotic dynamics must be confirmed with long simulation to ensure its stability.

**Figure 9:**
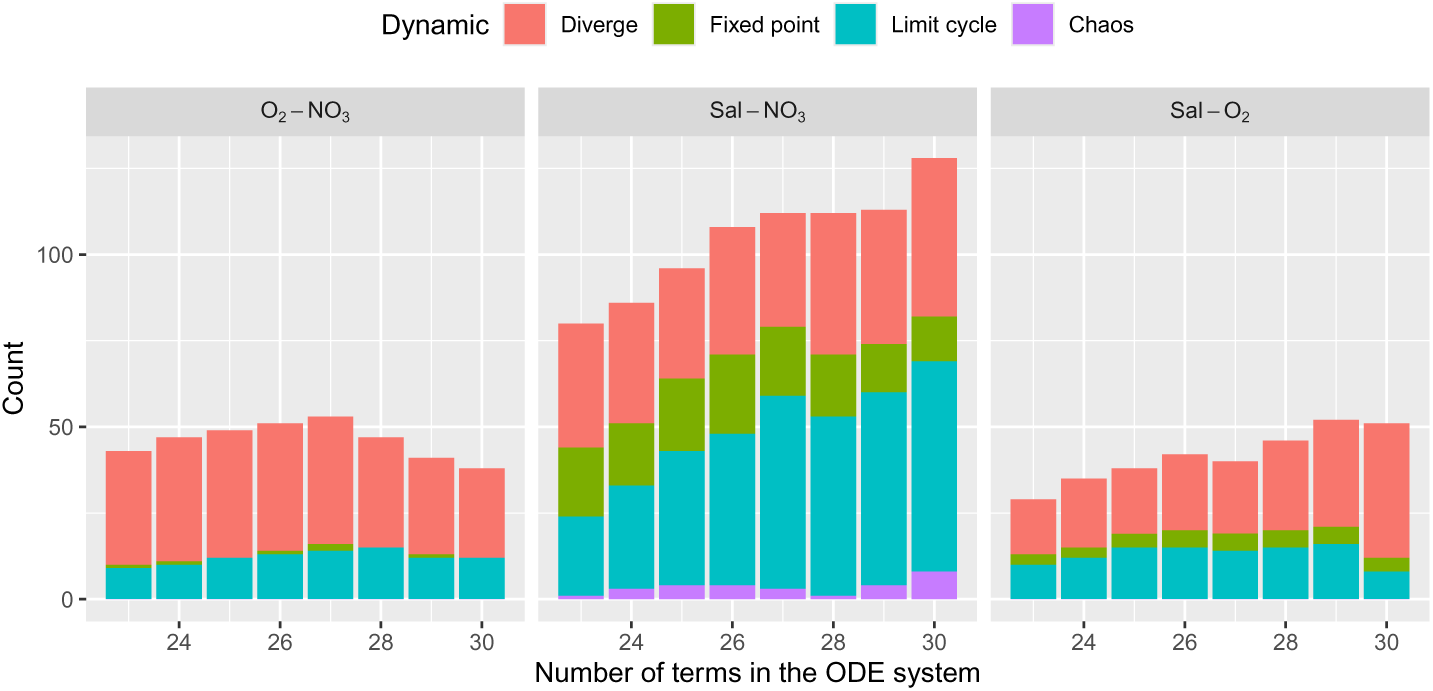
Dynamical analysis of the unclassified systems (range C, Fig. 3C). The numerical integration for a long integration time ensures the robustness of the classification. Over these 1600 ODE systems, our manual classification indicates that 50% diverge, 12% converge to a fixed point, 36% converge to a limit cycle and 2% converge to a chaotic attractor.

**Figure 10:**
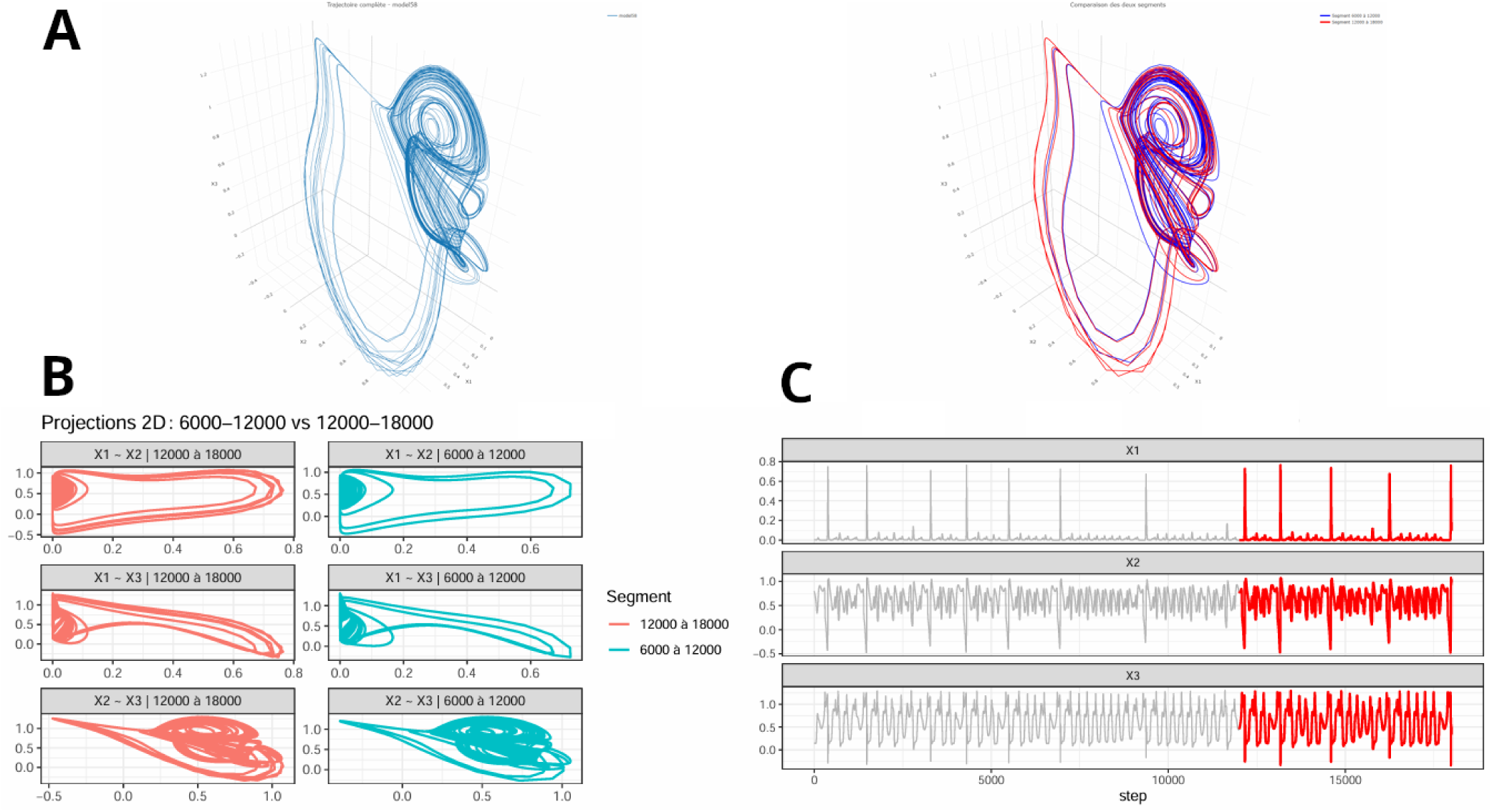
Automated plots for dynamical classification of a model generated via GPoM. Multiple outputs are obtained after a long numerical integration of a model performed with the function numicano in GPoM package (Mangiarotti, Le Jean, et al., 2023). Two colours are used to split data: integration from 6,000 to 12,000 points and the second from 12,000 to 18,000 points with a step equal to 0.01 in the numerical method. A. Three-dimensional phase spaces. B. Three couples of two variables in the phase space for assessing the globally time invariant property of chaotic attractors. C. Time series to detect anomalies. This visualisation permits to discriminate systems with transient chaos from chaotic dynamics.

#### B.1 Topological characterization

Including the entire dataset with the initial peak induces same results by generating more differential equation systems susceptible to describe the structure of the data with the couple salinity and NO_3_. However, only three attractors are solution to these systems (none including the latter couple of variables) with only two or three branches (Fig. 13). Templates are drawn using CATE (https://pypi.org/project/cate/ Olszewski et al., 2018) in Figs. 11, 13 and 14.

**Figure 11:**
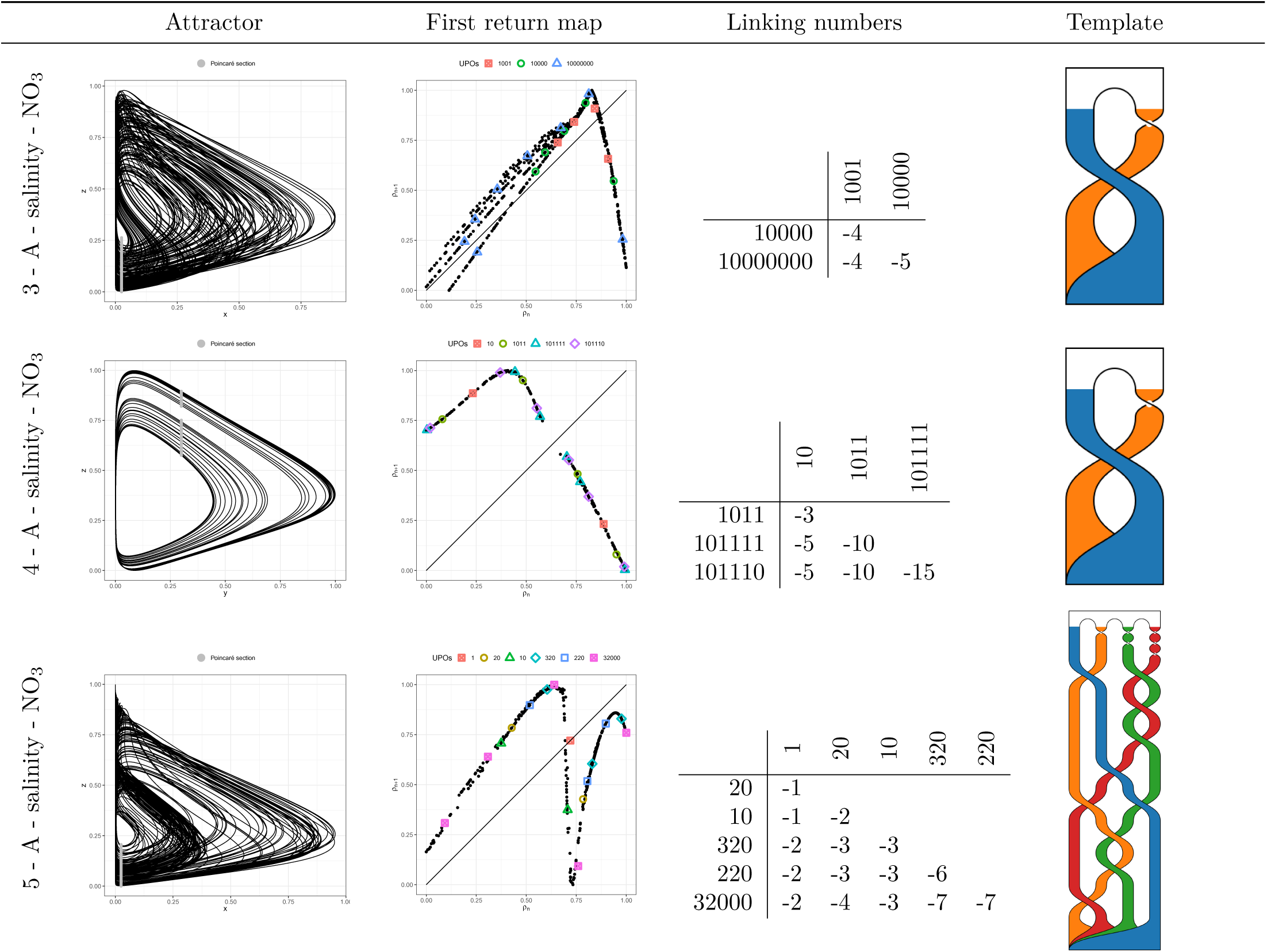
Examples of models having chaotic dynamics as solution under hypothesis Fig. 3A. Attractors variables values are normalized between 0 and 1.

In Fig. 13, only three attractors are solution of systems produced and none comes from the series that produces most systems (couple salinity - NO_3_).

**Figure 12:**
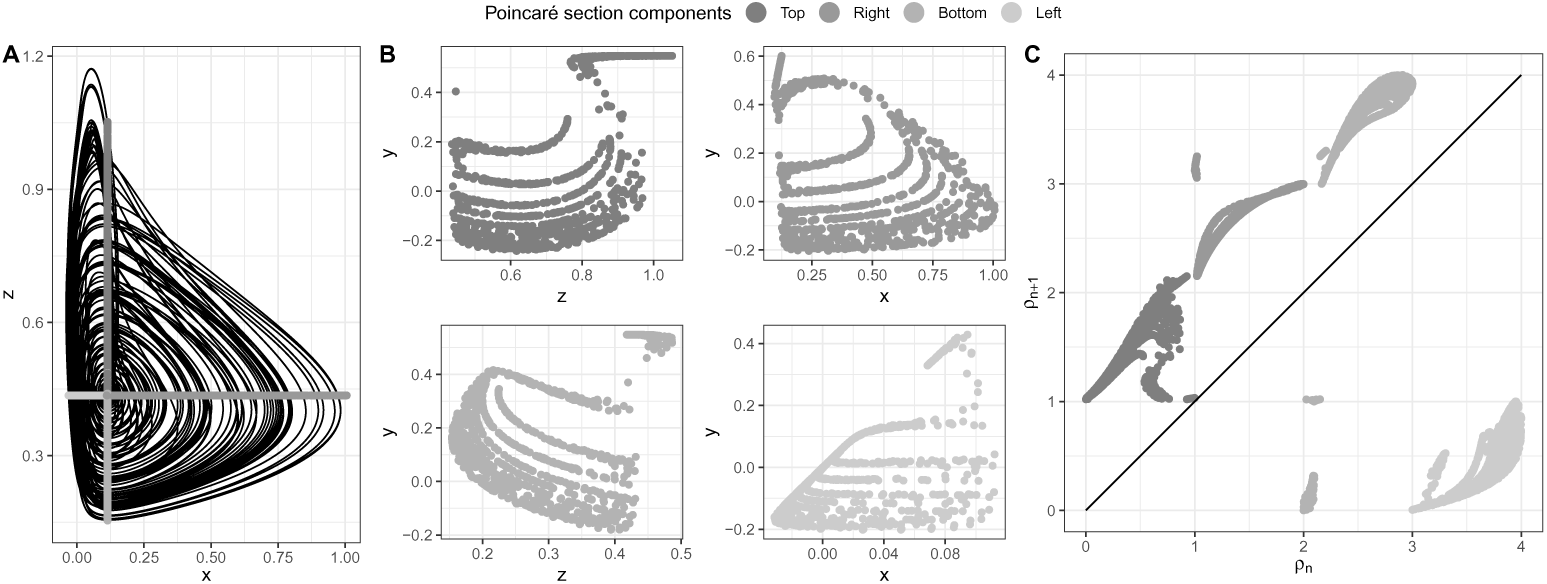
Complex attractor solution of model 9 from a weakly dissipative system with genus–3 bounded torus and a foliated structure. Using multiple components (A) to build a Poincaré section enables to details clockwise transitions between them with a first return map (C). The flow is not fully evolving clockwise because there are extra transitions from out of the clockwise scheme (top to right to bottom to left to top). Additional information required to obtain the topological characterisation of this attractor (see Rosalie, 2016; Rosalie and Letellier, 2015 for examples of genus–3 bounded templates and multiple components first return maps).

**Figure 13:**
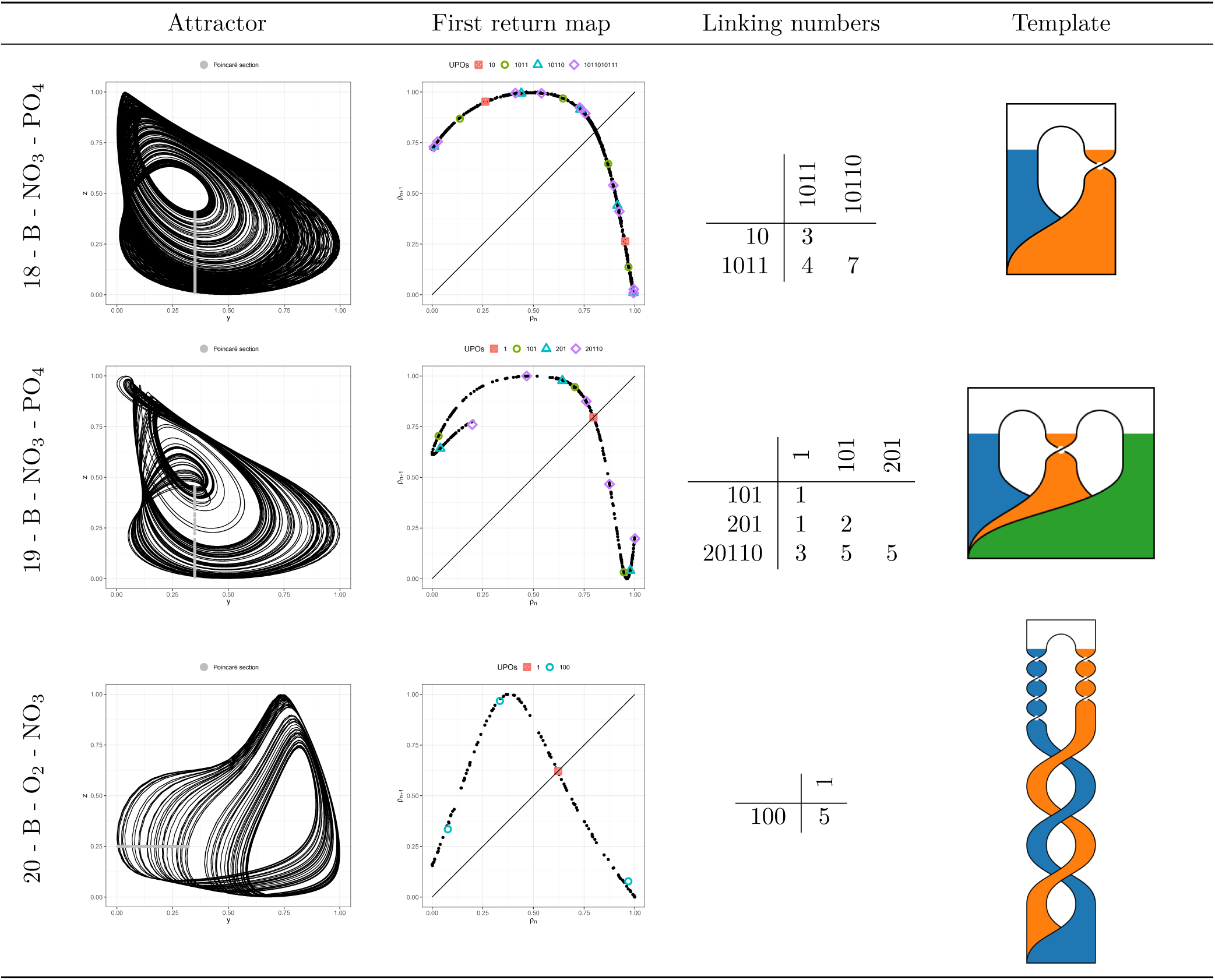
Hypothesis B including the entire dataset for all variables except temperature. Attractors variables values are normalized between 0 and 1.

**Table 3:**
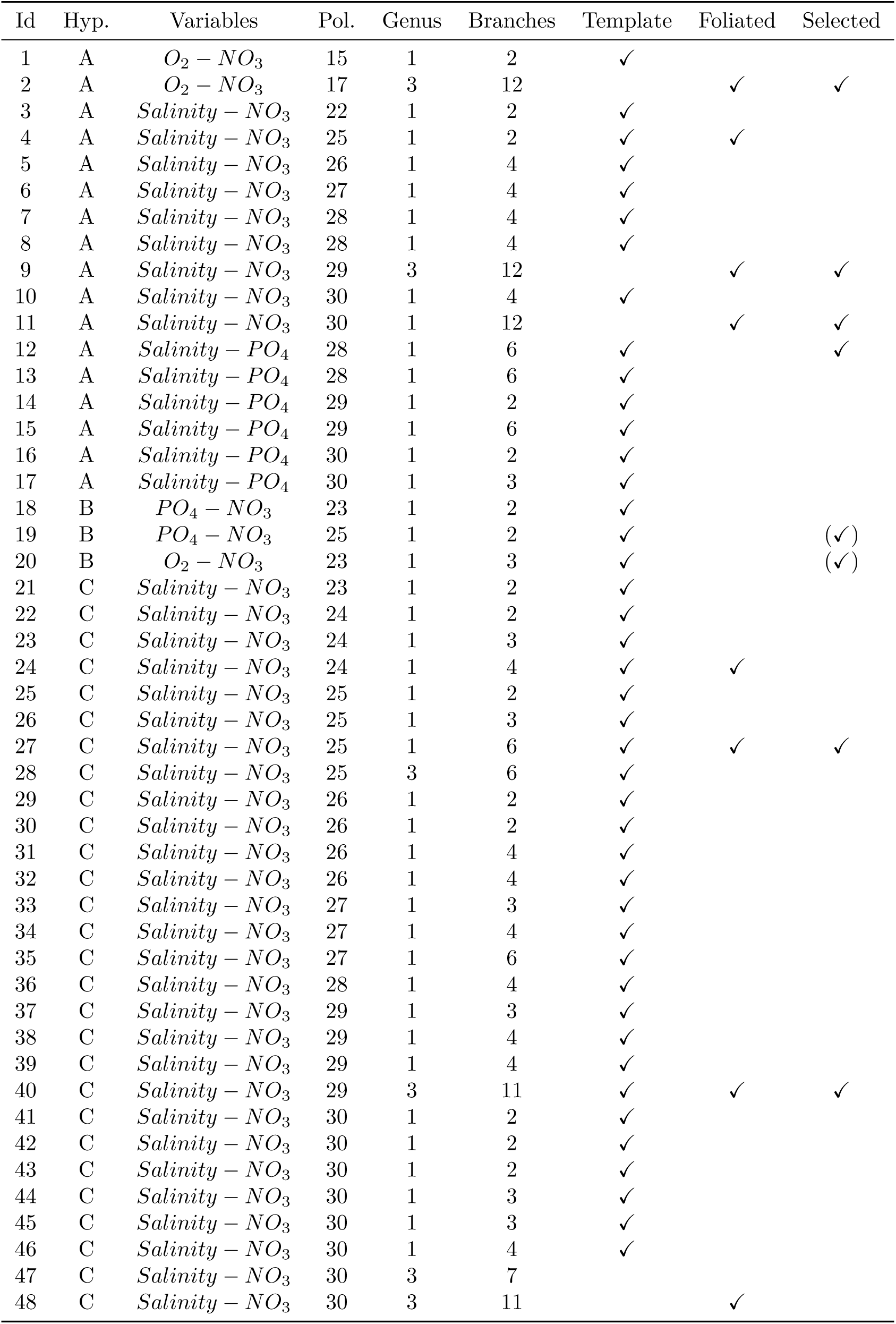
Pruning of models using dynamical properties. Models 19 and 20 are supposed to be selected but their range of values are far from the range of observed values to be used in comparison (Tab. 4).

**Table 4:**
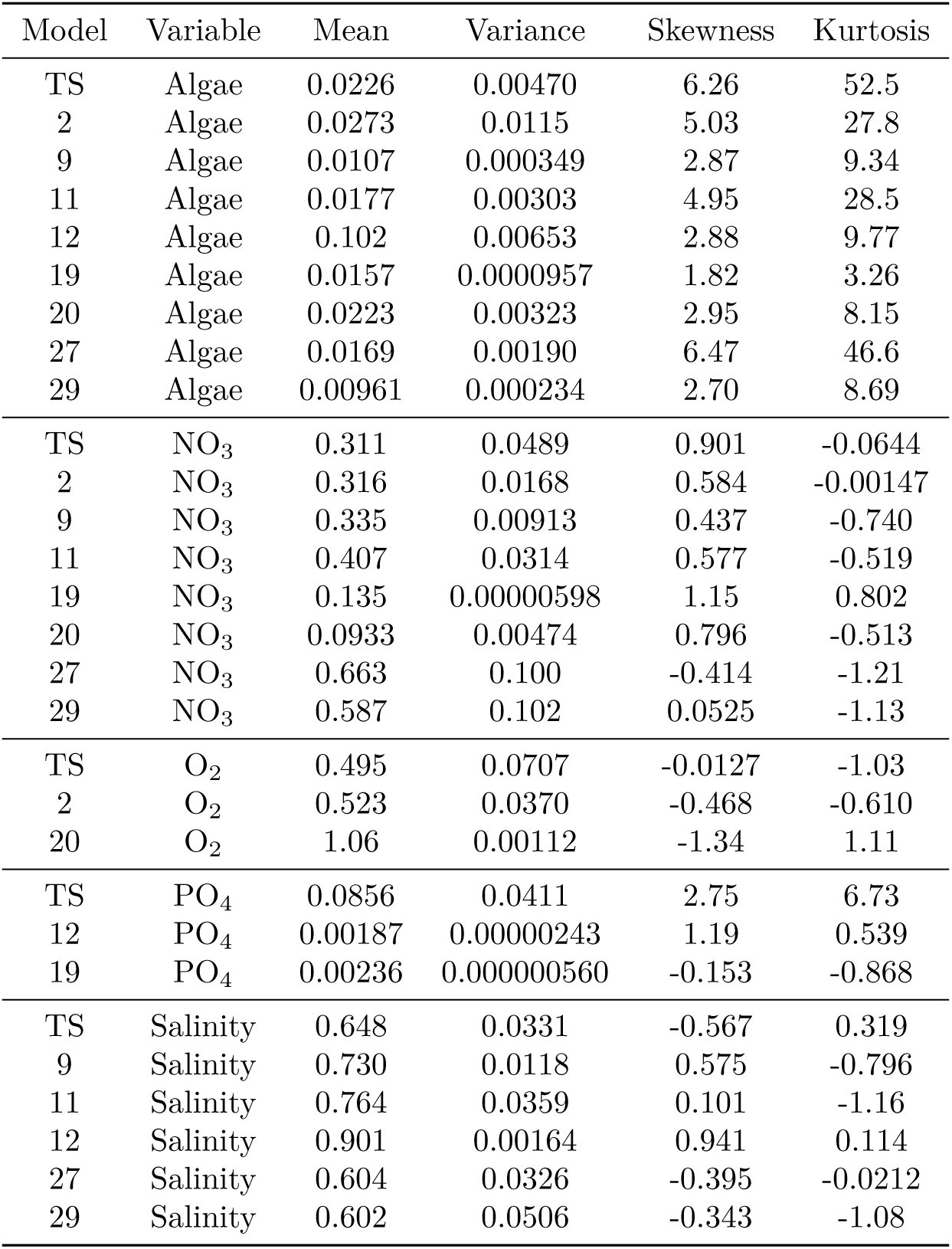
Statistical moments (mean, variance, skewness, kurtosis) for observations and model simulation. TS corresponds to the whole time series and values to list of models Tab. 3.

**Table 5:** Statistical moments (mean, variance, skewness, kurtosis) for attractor with distinct dynamics. Id of models is reported in Tab. 3. Model 47 is bounded by a genus–3 torus with complex dynamics while model 21 is a classical horseshoe mechanism of a chaotic attractor bounded by a genus–1 torus.

| Model | Variable | Mean | Variance | Skewness | Kurtosis |
| --- | --- | --- | --- | --- | --- |
| 21 | Algae | 0.00902 | 0.000237 | 1.88 | 2.60 |
| 47 | Algae | 0.00685 | 0.0000737 | 1.11 | 0.134 |
| 21 | NO<sub>3</sub> | 0.662 | 0.0509 | -0.205 | -1.47 |
| 47 | NO<sub>3</sub> | 0.778 | 0.0623 | -0.445 | -0.80 |
| 21 | Salinity | 0.535 | 0.0180 | 0.172 | -0.652 |
| 47 | Salinity | 0.603 | 0.0370 | 0.0804 | -1.10 |

**Figure 14:**
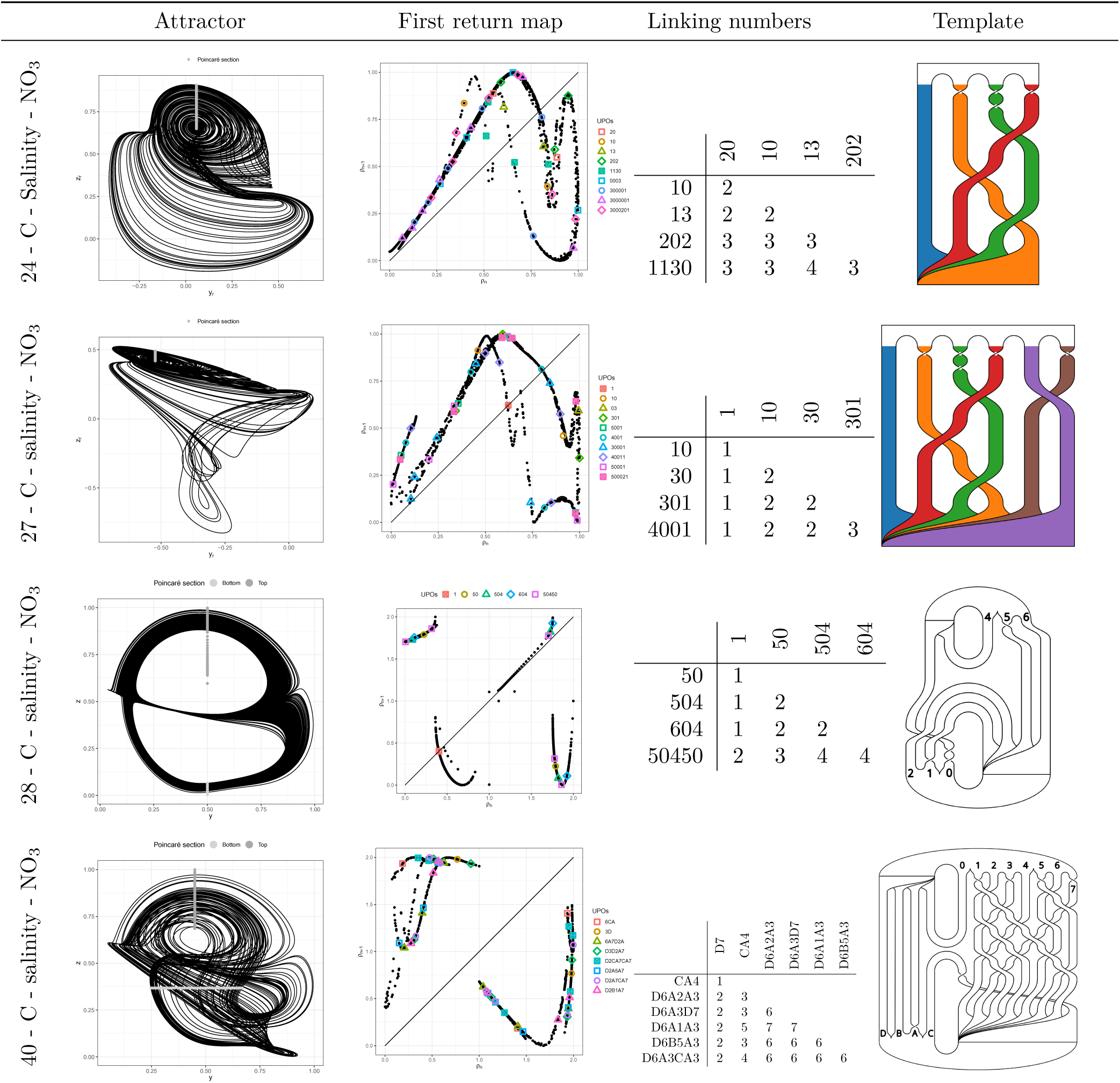
Topological characterisation of attractors under hypothesis. **C.** Attractors variables values are normalized between 0 and 1.

